# The genetic diversity of *Strongyloides papillosus* in Pakistani goats revealed by whole genome sequencing

**DOI:** 10.1101/2024.04.16.589736

**Authors:** Kiran Afshan, Yuchen Liu, Mark Viney

## Abstract

*Strongyloides* nematodes are parasites of livestock, and *S. papillosus* infects ruminant livestock that can cause disease. Recent genomic analysis of several *Strongyloides* species is now facilitating the population genomic analyses of natural *Strongyloides* infections, for example finding that *S. ratti* in wild UK rats exists as an assemblage of long-lived, asexual lineages. Here we have initiated an investigation into the population genomics of *S. papillosus* in goats in Pakistan. We find that *S. papillosus* is common, with a prevalence of 28%; that the population is genetically diverse and that individual goats commonly have mixed-genotype infections; and that there is evidence of only limited admixture. These results now provoke further questions about the host range of different *S. papillosus* genotypes that can in the future be investigated by further population genomic analyses.

## Background

*Strongyloides* is a genus of gut parasitic nematode that infects a wide variety of terrestrial vertebrates. *Strongyloides papillosus* has been reported to parasitise sheep, goats, and cattle and can be experimentally maintained in rabbits (Pinn et al., 2022). However, the species identity of *Strongyloides* infecting these host species has been questioned. Sequence analysis of the rRNA coding gene of *Strongyloides* from cattle and sheep found two different forms that appeared to be substantially (though not completely) host specific (Eberhardt et al., 2008). The names *S. papillosus* and *S. vituli* for parasites from sheep and from cattle, respectively, were originally suggested by Brumpt 1921 (though not widely used), but reinitiated by Eberhardt et al., 2008, with further support for *S. vituli* infection in cattle (Ko et al., 2019). More generally, the host specificity of *Strongyloides* species remains unexplored (Al-Jawabreh et al., 2023). Most *Strongyloides* species have been described morphologically, often from only some stages of the life cycle, which makes it hard to not only define a *Strongyloides* species, but also its host range. However, population genetic analyses of *Strongyloides* has the potential to resolve such questions.

The parasitic phase of *S. papillosus* (in common with other species of *Strongyloides*) consists of adult female worms only that live in the mucosal epithelium of the small intestine and reproduce by mitotic parthenogenesis. Eggs are shed in the host faeces, ultimately resulting in infective, filariform third-stage larvae (iL3s). These iL3s can develop either, directly, via two larval moults, or, indirectly, as the progeny of a facultative adult dioecious free-living generation that reproduces sexually (Viney, 2017). Hosts become infected by percutaneous infection by iL3s (Turner et al., 1960). The parthenogenetic reproduction of the parasitic female worms means that the larvae passed in faeces are genetic copies of their mother that is resident in the host gut. The only opportunity for sexual reproduction is when free-living males and female worms are present (presumably in faeces or nearby soil). The factors that control the occurrence of this free-living adult generation are not fully understood, but in *S. ratti* in rats this varies among *S. ratti* genotypes, and is affected both by the temperature external to the host and by the host immune response (Viney & Lok 2007). From a population genetic perspective the facultative nature of sexual reproduction may mean that the population genetic structure of *Strongyloides* deviates from that predicted by Hardy-Weinberg expectations.

*S. papillosus* can be a common infection of ruminants, with a prevalence of up to 58% reported in lambs, though the prevalence differs widely geographically and among different livestock management systems (Thamsborg et al., 2017). *S. papillosus* can cause diarrhoea, dehydration, anorexia, anaemia and malnutrition, especially in young animals (Thamsborg et al., 2017). High intensity infections can cause serious strongyloidiasis, which can be fatal (Pinn et al., 2022). Juvenile animals are most susceptible to heavy infection upon first exposure, but rapidly develop acquired immunity to future *S papillosus* infections (Turner et al., 1960). *S. papillosus* can be treated with anthelminthic drugs such as macrocyclic lactones (*e*.*g*. Ivermectin) and albendazole (Thamsborg et al., 2017). Rabbits can also be infected with *S. papillosus* and as such can be used as an experimental model for *S. papillosus* (Thamsborg et al., 2017).

The genome of *S. papillosus* has been sequenced, together with 3 other species of *Strongyloides* (Hunt et al., 2016). The *S. papillosus* genome is estimated to be 60Mbp, though it is less well assembled than the other *Strongyloides* species such that the estimate of its genome size may not be very accurate. Notwithstanding, the *S. papillosus* genome assembly can be used for population genomic analyses of this species.

Population genetic analyses of parasitic nematodes of livestock has found that there is very high gene flow and an absence of parasite population structure (Blouin et al., 1995). This was thought to be due to both the often very large effective population sizes of the nematodes, and because of extensive host movement that occurs as part of commercial farming. For nematode parasites of species of wildlife then there are a diversity of observed parasite population structures, with this dependant on host and parasite specific biology (Cole and Viney 2018). There has been previous population genomic analysis of *Strongyloides*. Specifically, *S. ratti* infecting wild rats in the UK was found to exist as a mixture of mainly asexual lineages parasites that were widely dispersed across the host population, findings that were consistent with earlier, UK-wide, studies (Fisher & Viney, 1998; Cole et al., 2023).

Here we report an initial population genomic analysis of *S. papillosus* in goats in Pakistan where we have used single worm, whole genome sequencing. We describe the patterns of genetic diversity among these worms, find evidence of mixed-genotype infections in hosts, but limited admixture.

## Materials and Methods

### Study area, faecal sampling and culturing

This study was approved by the bioethical committee of Quaid-i-Azam University, Islamabad, Pakistan. Sampling was conducted at four slaughterhouses within Rawalpindi City, with the slaughterhouses ranging from a minimum of 9 to a maximum of 32km between them. Faecal samples were collected directly from the rectum of 150 slaughtered goats using sterile disposable plastic gloves, with fresh gloves used for each goat to avoid cross-contamination. The faeces were cultured by mixing a few grammes of fresh faeces with an approximately equal volume of wet charcoal, which was then kept at room temperature for up to 13 days when individual iL3s were collected, which were identified as *Strongyloides* by morphological examination (Zhou et al., 2019); no free-living adult males or females were observed.

### DNA extraction and genome sequencing of single larvae

The DNA extraction protocol is based on the method of Cole et al., 2023 but adapted for ethanol-stored worms after Zhou et al., 2019. Pools of larvae from each host were processed separately, individual larvae in TE were placed into wells of a 96 well plate; lysis buffer was added to give a final concentration of 200 mM NaCl,100 mM Tris-HCl (pH8.5), 50 mM EDTA (pH8), 0.5 % w/v SDS, 0.9 mg /mL Proteinase K, and 45 mM Dithiothreitol (DTT). Plates were then sealed and held at 60°C for 2 hours with gentle agitation, after which they were held 85°C for 15 minutes to inactivate the Proteinase K. Lysates were then stored at -80°C before whole genome sequencing.

Whole genome sequencing was performed by the Centre for Genomic Research, University of Liverpool. Illumina libraries were prepared using the NEB FS Ultra 2 DNA kit at half of the manufacturer’s specified reaction volumes.

### Sequence analyses

We whole genome sequenced 114 *S. papillosus* iL3s, mapped the sequence reads using Bowtie2 to the *S. papillosus* reference genome (PRJEB525, WBPS18 available from WormBase-ParasiteSite; parasite.wormbase.org), and calculated the average depth and coverage across the genome of each alignment using SAMtools. Of these 114 larvae, 37 had a genome coverage of at least 70% and an average depth of at least 10, and these samples were then analysed further.

Single Nucleotide Polymorphism (SNP) calling was performed using SAMtools. We used VCFtools to filter SNPs using the criteria that (i) they had a mean mapping quality of at least 20, (ii) they had a QUAL score of at least 20, (iii) SNP loci had a minor allele frequency of ≥ 0.02, and (iv) their mean depth was between the 10 and 90% percentiles of the whole data set, which was 6 and 21, respectively. Among the 37 *S. papillosus* genome sequences, nucleotides that were identical among all samples (but different from the PRJEB525 reference genome) were removed.

Using the filtered SNP set of the 37 *S. papillosus* iL3s we performed population structure analysis by calculating the pairwise genetic similarity among the 37 worms and then (i) using principal component analysis (PCA) in PLINK 1.9 and Tassel v.5 with plots visualized in R studio v.4.2.1; (ii) constructing a neighbour-joining tree in Tassel, visualising the tree in iTOL; and (iii) using ADMIXTURE version 1.3.0 (Alexander and Large, 2011) analysis, with linkage disequilibrium (LD)-based pruning, which was done with window sizes of 50 kb, step sizes of 10 SNPs, and with a r^2^ threshold of 0.1. ADMIXTURE was run for k = 2 – 15 and results plotted in R studio v.4.2.1. To investigate the genetic distance among worms we calculated the identity-by-distance (IBD) on the filtered SNP set in PLINK. IBD values were categorised into two pairwise comparison groups (i) from within the same host and (ii) and among different hosts.

The *S. papillosus* reference mitochondrial genome was obtained from NCBI GenBank, NC_028622.1. We generated maximum likelihood trees for the mitochondrial genomes of the 37 larvae, producing consensus fasta sequences for each; where an individual was heterozygous the reference allele was applied. Sequences were aligned with MAFFT (Katoh et al., 2009) using the fast alignment method FF-NST-1. Maximum likelihood tree estimation was carried out utilizing RaxML (Stamatakis, 2006), applying a general time reversible gamma model of substitution rate heterogeneity, and rapid bootstrapping with 100 replicates and visualized in ITOL.

It has been suggested that there are two species of *Strongyloides* that infect ruminants, *S. papillosus* and *S. vituli*, which can be differentiated by their rRNA sequence (Eberhardt et al., 2008; Ko et al., 2019), and a complete mitochondrial genome sequence has also been reported for *S. vituli* (Ko et al., 2023). For the 37 samples we analysed the rRNA and mitochondrial sequences to confirm their species identity.

## Results

We sampled 150 goats of which 42 (28%) were faecal culture positive for *Strongyloides*. In all we obtained 526 iL3s, ranging from 7 – 15 iL3s per infected goat. We sequenced 114 individual iL3s of which 37 generated sufficiently good data and which were taken forward for further analysis. These 37 larvae were obtained from 21 goats. For these 37 this resulted in sequence data with an average genome coverage of 89% (range 74 – 94%), and an average read depth of 21 (range 9.7 – 69), giving an average genome coverage size of 53.6 Mb. We detected 11,023 filtered SNPs in the nuclear genome, giving an average SNP density of 0. 2 SNPs per 10 kb. This alignment of reads to the *S. papillosus* genome is consistent with identifying these larvae as *S. papillosus* rather than *S. vituli*. We also analysed the rRNA and mitochondrial genome sequence of the 37 samples, which confirmed their identity as *S. papillosus*.

To investigate the genetic relationship among the 37 iL3s we calculated the pairwise nuclear genetic similarity among the larvae and used this to construct a neighbour joining tree (**Figure 1A**). The shape of this tree showed that there was extensive genetic heterogeneity among the worms. In this tree most (33 worms) were on short-length branches, showing their relatively close genetic similarity. But two pairs of worms (two of which were from a single host, the other two from two different hosts) were on long, distant branches in the tree, showing their relative genetic dissimilarity from the other 33 worms. A PCA of the same data also showed three clusters of iL3s, with 33 iL3s in a single cluster, and then two smaller clusters each consisting of the same two pairs of divergent iL3s (**Supplementary Figure 1**). We investigated the data for evidence of admixture, with the results for k = 10 (Cross Validation (CV) = 0.13) showing that only eight worms (22%) had evidence of admixture (**Figure 1B**). Examining admixture results for other values of k, also showed low levels of admixture; specifically for k = 3, 7, 15 then six, three, and six worms showed admixture, respectively (**Supplementary Figure 2**)

**Figure 1.**
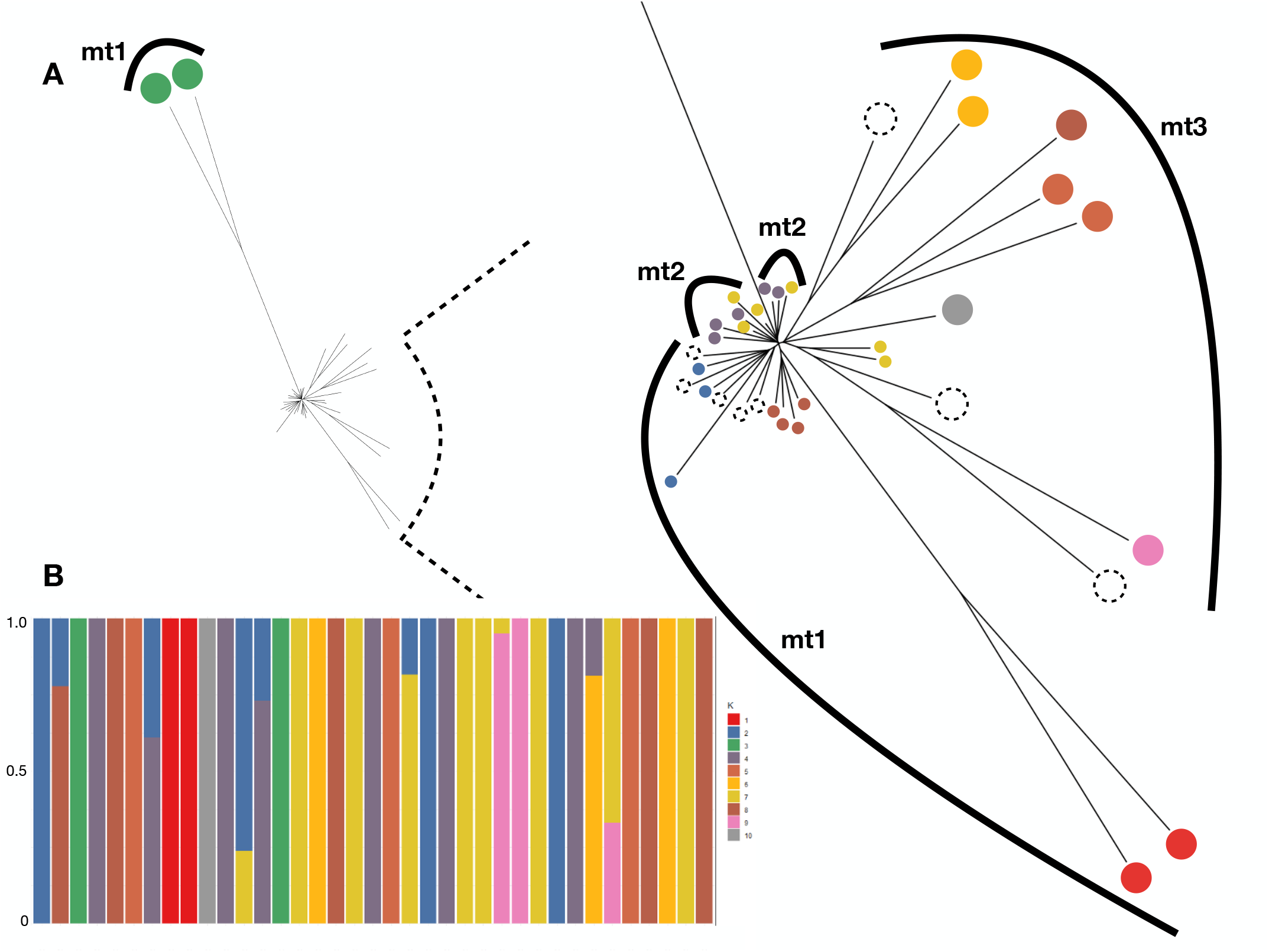
Population genomic analyses of 37 *S. papillosus* larvae from 21 goat hosts. (A) Neighbour-joining tree of the nuclear genome (full view to the left, and the central region enlarged to the right), with individual worms coloured according to the ADMIXTURE groups, with the eight admixtured worms shown as dotted circles and the mitochondrial clades shown as numbered (mt1, mt2, mt3) arcs and (B) ADMIXTURE analysis showing the degree of ancestry on the y-axis, with K = 10. In (A), the two worms from ADMIXTURE group 3 (green) are from different hosts; two worms in ADMIXTURE group 1 (red) are from the same host.

The mitotic parthenogenetic reproduction of *Strongyloides* parasitic females means that offspring of a single parasitic female worm will be genetically identical, and so that genetically identical iL3s would be observed in host faeces. Conversely, if hosts are infected with genetically different parasitic females, then genetically diverse iL3s will be present in hosts faeces. With this in mind, we considered hosts from whom multiple iL3s were sequenced. Among the 21 hosts, 11 each had more than one worm sequenced, giving a total of 27 worms (five hosts with three worms; six hosts with two worms). We compared the average genetic distance of worms from within hosts, with worms from among different hosts, finding that there was little difference: mean (SD) within hosts 0.063 (0.068), but 0.059 (0.059) among hosts. This suggests that hosts may harbour genetically different worms. We asked whether worms from a single host were in one or more k = 10 ADMIXTURE-defined groups and / or were admixtured. We found that 10 of the 11 hosts contained worms from more than one ADMIXTURE-defined group and / or were admixtured. This suggests that multi-genotype *S. papillosus* infections are common in goats in Pakistan.

For the mitochondrial genome we detected 357 SNPs and used these to construct a maximum likelihood tree of the 37 iL3 and the reference *S. papillosus* mitochondrial genome. This showed that there were three *S. papillosus* clades (**Figure 2**). Two clades contained most (16 and 9 iL3s ) iL3s, with the third clade containing 12 iL3s that were more divergent. This third clade also contained the *S. papillosus* reference mitochondrial genome (**Figure 2**).

**Figure 2.**
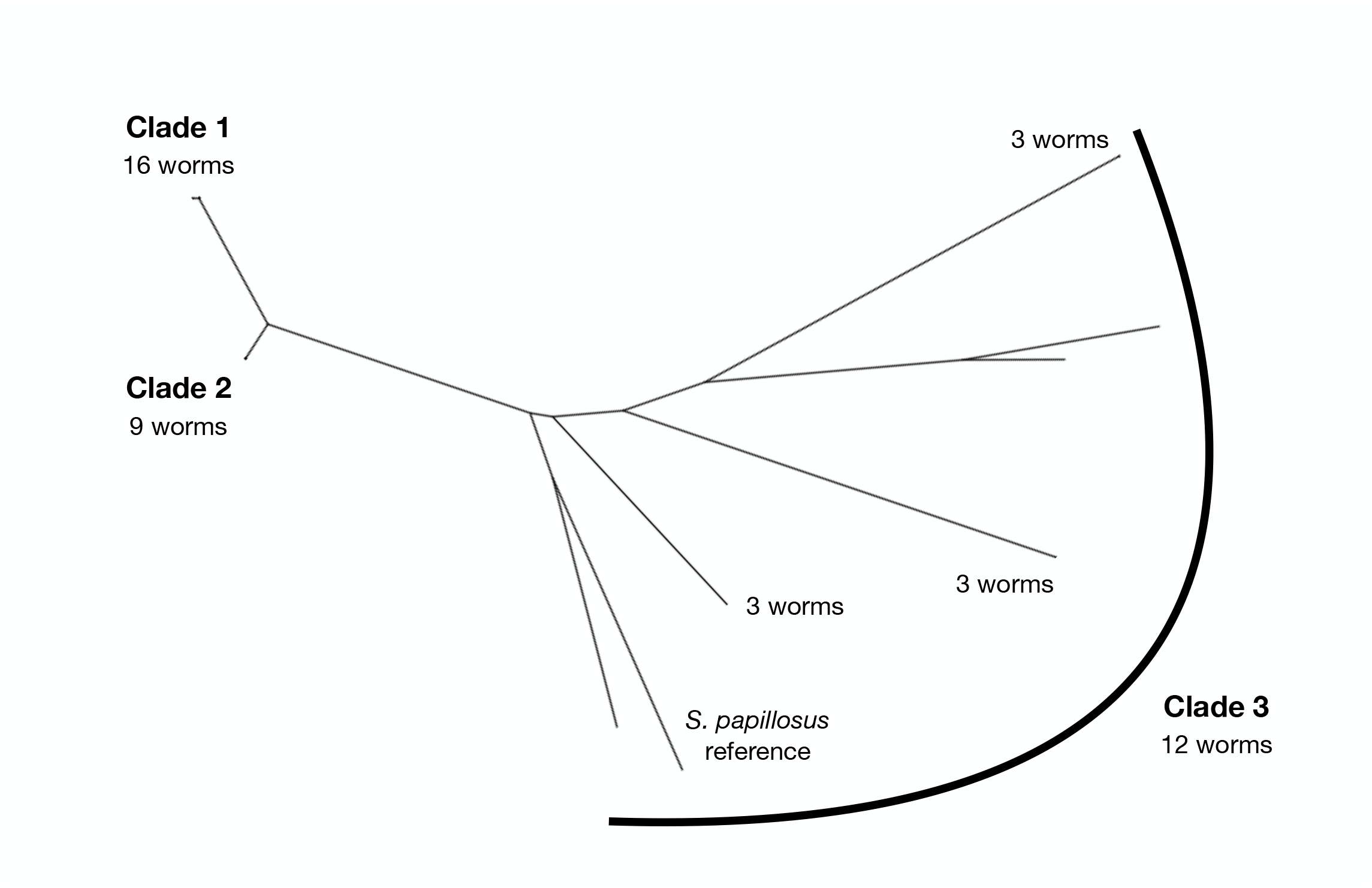
A maximum likelihood tree of the mitochondrial genome of the 37 samples and the *S. papillosus* reference mitochondrial genome.

As for the nuclear genome, we considered the 11 hosts that each had more than one sequenced worm. We found that nine hosts contained worms that were in more than one mitochondrial clade. This is further evidence of mixed *S. papillosus* genotype infections in these hosts. Interestingly, the four iL3s with divergent nuclear genomes (**Figure 1A**) were within mitochondrial clade 1 (**Figure 2**). This result therefore shows some divergence in the genetic relationship among iL3s as judged by nuclear and mitochondrial data.

We compared the nuclear and mitochondrial trees. This showed that, with the exception of one nuclear clade, the three mitochondrial clades grouped related nuclear clades (**Figure 1A**). The exception is the highly divergent nuclear clade, which belongs to mitochondrial clade 1, whereas its closest neighbours belong to mitochondrial clade 2.

## Discussion

This work has used whole genome sequencing to describe the genetic diversity of *S. papillosus* in goats in Pakistan. We have found that *S. papillosus* is a common infection of goats in Pakistan, with a prevalence of 28%. *S. papillosus* can infect a number of species of ruminants, and it remains to be discovered what its prevalence is in these other species. Given that *S. papillosus* is able to infect a number of different potential host species, there is the possibility for among-host species transmission of *S. papillosus*. Such sharing of parasites among different host species has the potential to genetically structure the parasite population, with this possibility being dependant on the amount of within, verses between, host species transmission.

This work revealed evidence of extensive genetic diversity among the parasites sampled, for both the nuclear and mitochondrial genomes. The mitotic parthenogenetic reproduction of *Strongyloides* parasitic females means that their offspring will be genetically identical, though sequencing errors will not result in identical gene sequences. Our data show that mixed-genotype infections are common in the goats that were sampled, which is likely due to goats being repeatedly exposed to infection from a genetically diverse *S. papillosus* population. This pattern is broadly consistent with patterns of population genetics seen in other parasitic nematodes (Blouin et al., 1995). Among the parasites sampled, a minority were genetically relatively more divergent. Specifically, by analysis of the nuclear genome 4 of the 37 worms were more distinct. This might point to the existence of even higher levels of genetic diversity in the *S. papillosus* population, which might be revealed by more extensive sampling in goats. Further, because *S. papillosus* is able to infect different ruminant species, the more-divergent genotypes we observed may be because these are parasites whose host preference (but not restriction) is for species other than goats. However, further sampling and genetic analysis is needed to understand the distribution of *S. papillosus* among different ruminant host species and the relationship of *S. papillosus* and *S. vituli* (Eberhardt et al., 2008). The results of the analysis of the mitochondrial genome are broadly consistent with the analysis of the nuclear genome, showing 3 distinct clades with the third clade containing mitochondrially diverse worms (and the *S. papillosus* reference genome). Again, these results may potentially indicate a diversity of parasites that infect a range of host species, of which only a small portion has been sampled in the current study. However, it is notable that the grouping of parasites by their nuclear and mitochondrial genomes is not fully concordant. This is indicative of different histories of these two genomes in some of the parasites that we have sampled, which has also been observed in *Ascaris* (Anderson et al., 1997).

The *Strongyloides* life cycle is obligatorily parthenogenetic with facultative sexual reproduction, where the degree of sexual reproduction varies between species, geographical location, and genotype (Viney & Lok 2007). The amount of sexual reproduction in *S. papillosus* that we have sampled is unknown, but during the culture of faeces we did not observe any free-living adult stages, suggestive of rare or no sexual reproduction. A previous analysis of wild *S. ratti* in the UK found that it consisted of a mixture of long-lived asexual lineages (Cole et al., 2023), and a similar pattern may exist among the *S. papillosus* that we have sampled, though further study is required to confirm this.

To extend and develop this work and to better understand the population genomics of *S. papillosus* it would be desirable to more extensively sample larvae from a diversity of sympatric host ruminants. This will allow an analysis of the degree of partitioning, if any, of *S. papillosus* genetic diversity among different host species. Future work would also be facilitated by a better, more complete assembly of the *S. papillosus* genome, which could be achieved by long-read DNA sequencing. Analysis of diverse, wild genotypes of *S. ratti* has discovered clusters of parasitism genes (Hunt et al., 2016) that are hyperdiverse compared with the rest of the *S. ratti* genome (Cole et al., 2023). It would be interesting to study this in *S. papillosus* too, but this will also require a more complete genome assembly.

## Acknowledgments

This work was funded by an International Veterinary Vaccinology Network Laboratory Exchange Award and Quaid-i-Azam University, Islamabad Pakistan. YL is supported by the University of Liverpool and the China Scholarship Council.

## Supplementary Figures

### Supplementary Material

**Supplementary Figure 1.**
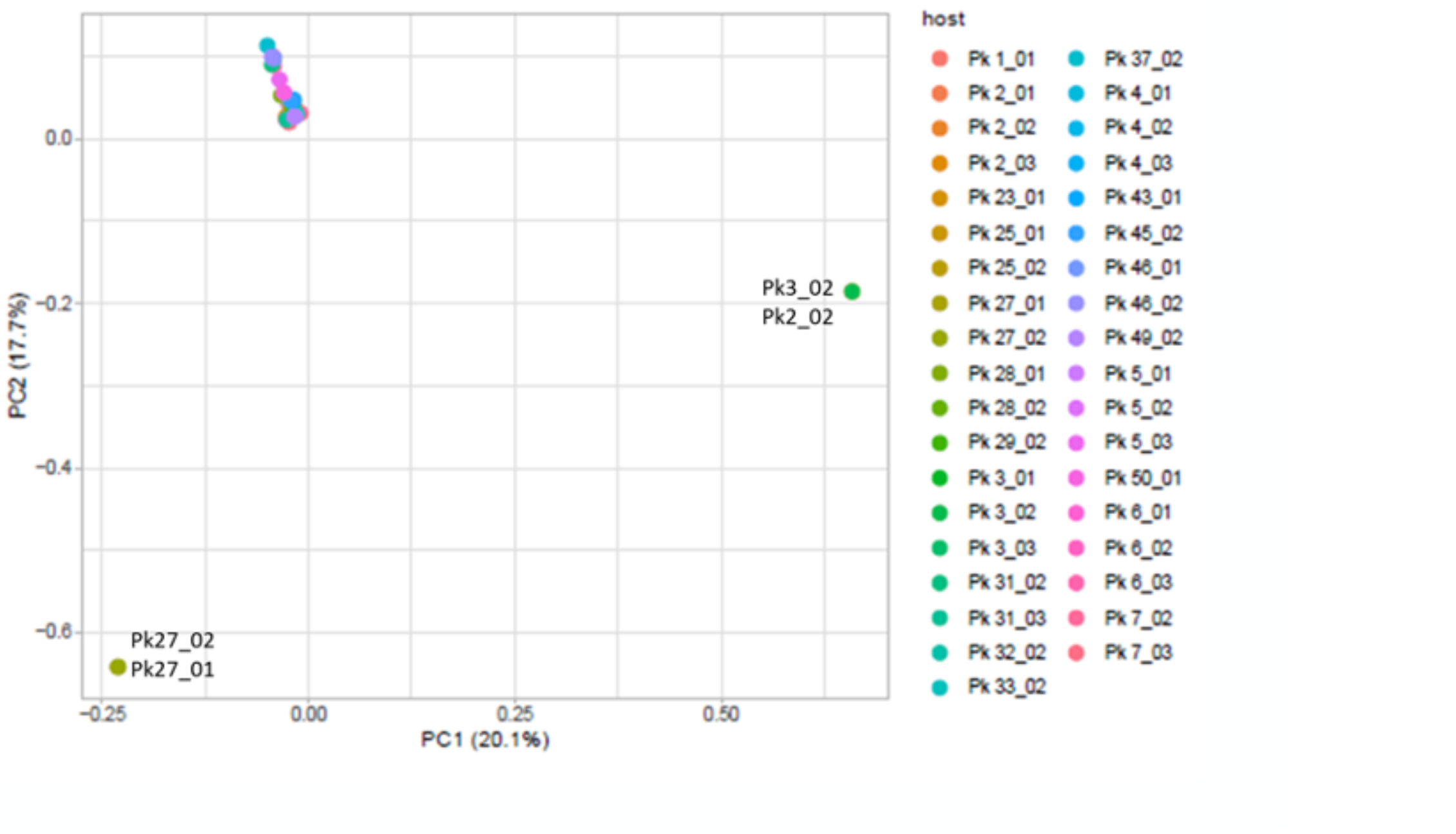
Principal component analysis showing the first two principal components, which accounts for 38% of the variance.

**Supplementary Figure 2.**
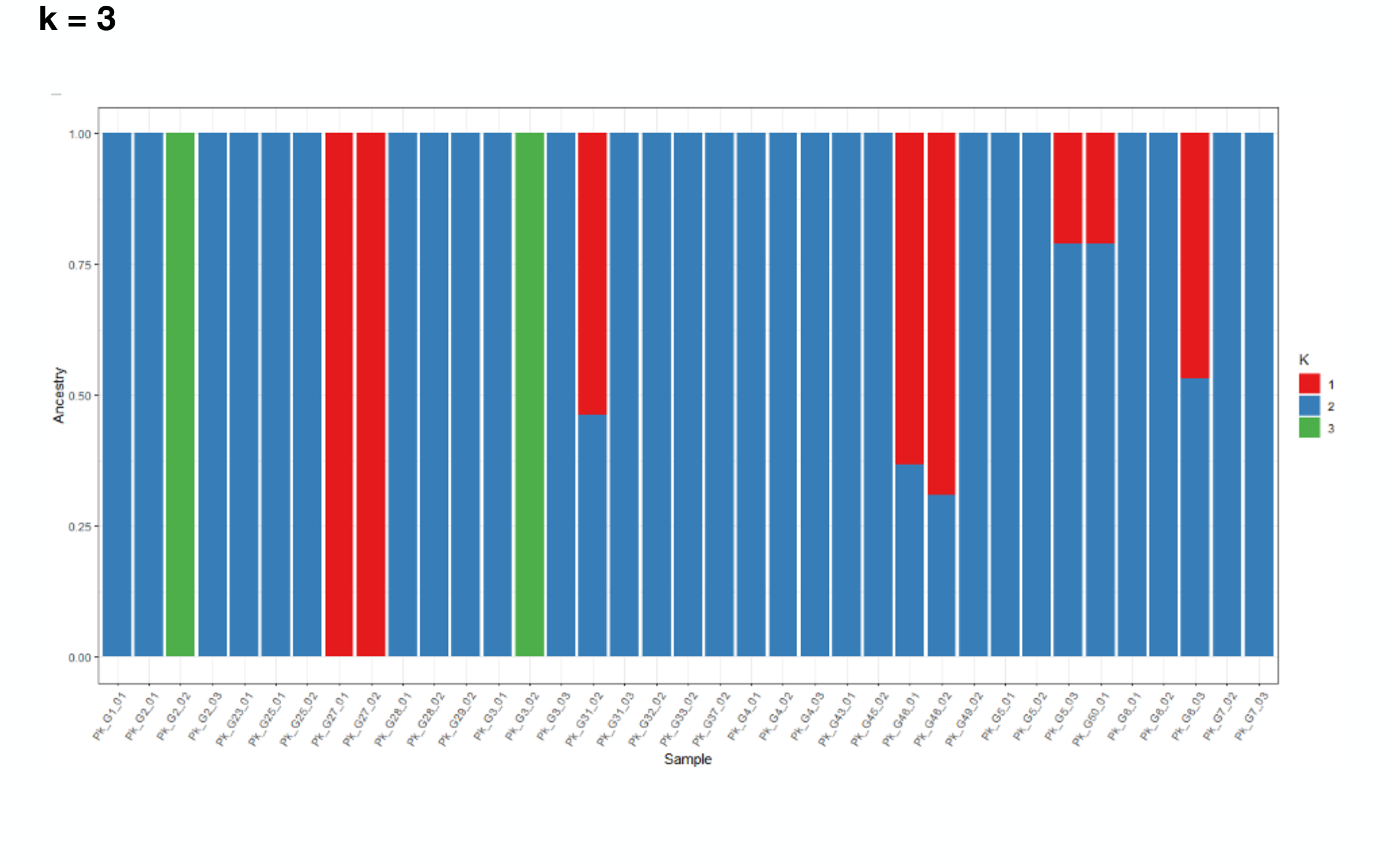
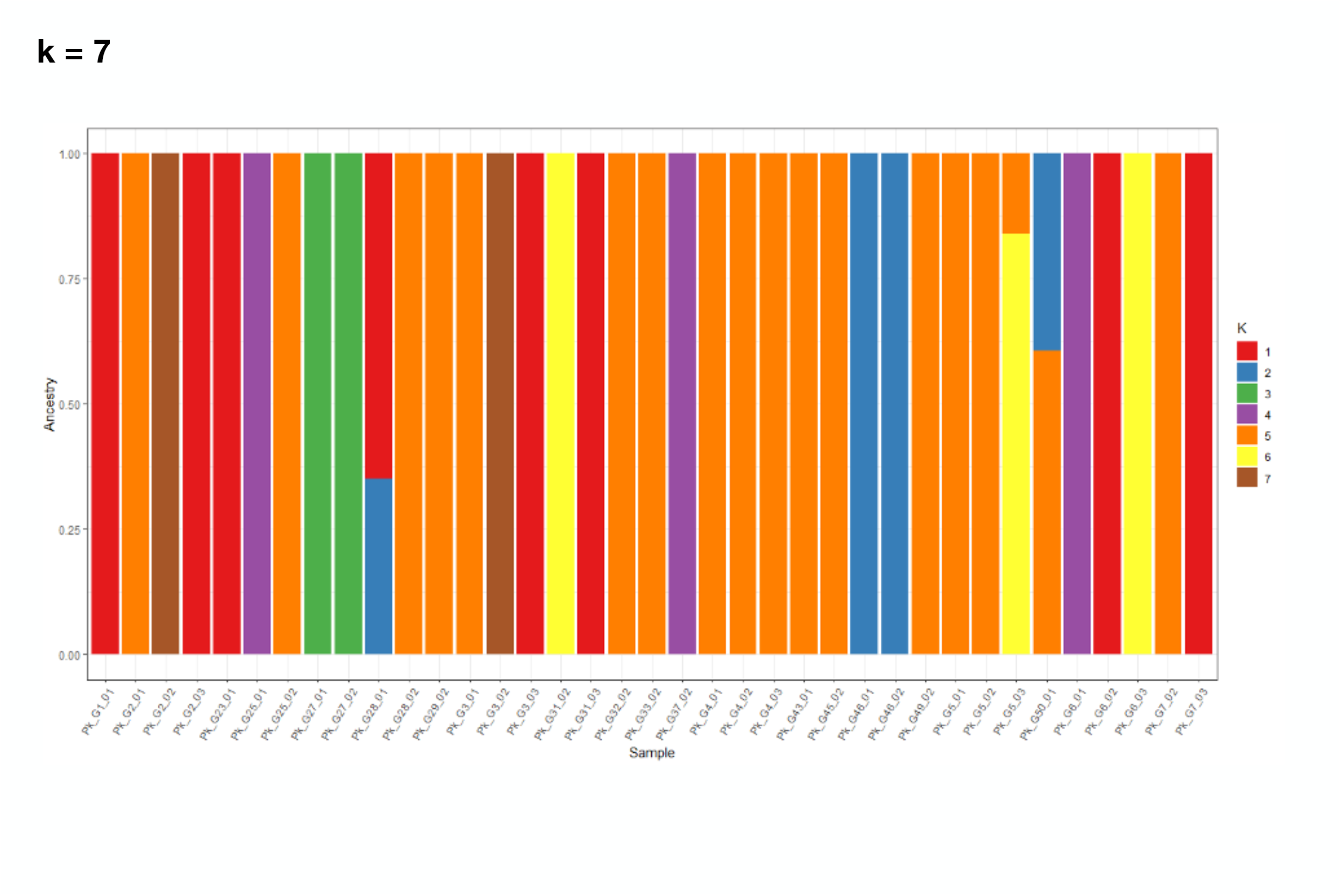
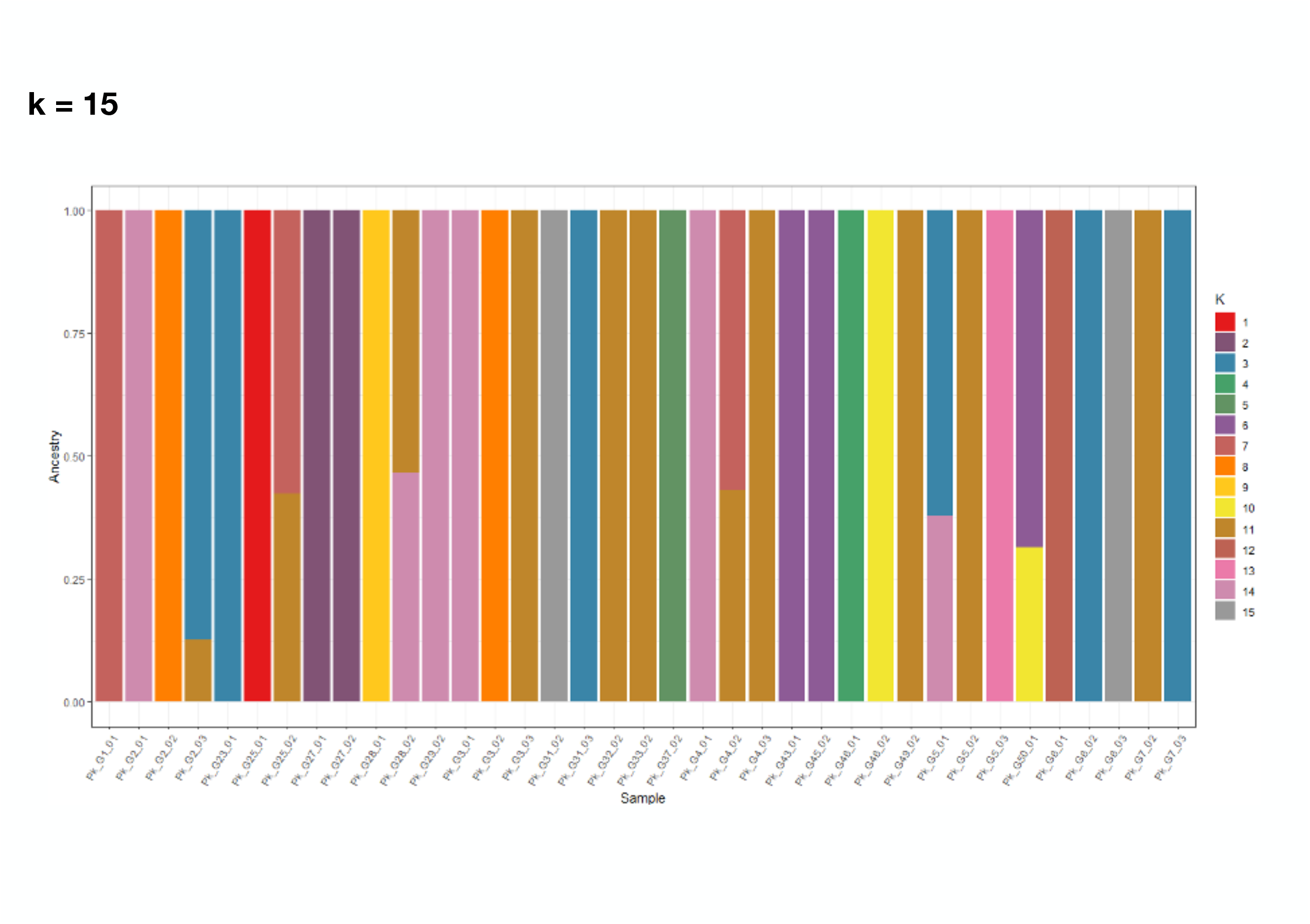
ADMIXTURE analyses for k = 3, 7 and 15.

